# Mesophyll-Specific Circadian Dynamics of CAM Induction in the Ice Plant Unveiled by Single-Cell Transcriptomics

**DOI:** 10.1101/2024.01.05.574430

**Authors:** Noé Perron, Christopher Dervinis, Wendell Pereira, Brad Barbazuk, Matias Kirst

## Abstract

Crassulacean acid metabolism (CAM) is an evolutionary modification of the C_3_ photosynthetic carbon dioxide fixation pathway used by approximately 7% of terrestrial plants to live in drought-prone environments. Facultative CAM species, such as *Mesembryanthemum crystallinum* (common ice plant), possess the unique ability to switch from C_3_ to CAM photosynthesis in response to high-salinity and water-deficit stress. Here we characterized the environmentally-triggered transition from C_3_ to CAM in the ice plant using single nucleus RNA sequencing (snRNA-seq) to identify its putative regulators, supported by a novel high-quality assembled and annotated genome. Analysis of snRNA-seq datasets from ice plant leaves transitioning between C_3_ and CAM collected at dawn and dusk revealed substantial transcriptional changes in mesophyll cells at the onset of CAM induction. Notably, our findings identify mesophyll sub-cell types engaged in either CAM or C_3_ photosynthesis at dusk. Cell trajectory inference analysis reconstructed both 24-hour CAM and C_3_ cycles, enabling a direct comparison of gene expression profiles in these pathways. This comparative study uncovered divergent expression patterns of key circadian clock genes in CAM and C_3_ cell trajectories, pointing to a connection between circadian regulation and CAM induction.

## INTRODUCTION

The ancestral C_3_ photosynthetic carbon assimilation pathway is estimated to occur in 90% of land plants^1,2^. The adaptation to varying environmental conditions, including periods of low atmospheric CO_2_, scarce water supply, and increased evaporative demand, has driven the evolution of alternate carbon fixation mechanisms^2^. C_4_ photosynthesis has evolved in approximately 3% of flowering plant species, and Crassulacean Acid Metabolism (CAM) is present in roughly 7%^1,2^. CAM holds potential economic and environmental value due to its unique mechanism of fixing CO_2_ at night. This process involves nocturnal stomatal opening, leading to lower evapotranspiration rates and higher water retention in leaves compared to C_3_ and C_4_ plants that fix CO_2_ during the day^1,3–6^. CAM plants demonstrate remarkable drought tolerance, exhibiting up to 20 times greater water-use efficiency than C_3_ plants^6^. Unlike obligate CAM species, which consistently exhibit CAM in fully-developed tissues, facultative CAM plants, such as the common ice plant (*Mesembryanthemum crystallinum*), can switch from C_3_ to CAM under environmental stresses like drought^3,5,7–9^. This transition has been prominently studied in the ice plant, making it a model species for understanding facultative CAM and discovering its regulators^3,7,10,11^. However, engineering CAM into non-CAM plants is still challenging due to incomplete knowledge of its regulation throughout the 24-hour cycle.

CAM operates in a 4-phase diel cycle model^12,13^. Phase I involves CO_2_ assimilation and carboxylation by phosphoenolpyruvate carboxylase (PPC), leading to malic acid accumulation in the vacuole and a significant reduction of cellular pH^5^. Phase II, spans the early part of the light period. Phase III, the day phase, is characterized by the decarboxylation of C_4_ acids and CO_2_ integration into the Calvin-Benson-Bassham (CBB) cycle. Phase IV marks the end of the light period. While the circadian regulation of phase I is well-described in several CAM species, the mechanisms regulating phases II and IV remain largely unknown.

In plants, circadian clock control is not only temporally, but also spatially organized^14^. Unlike the mammalian clock, which is a centralized process, the plant clock has been reported to act in a tissue and cell type-specific manner^14^. This study explores the C_3_-to-CAM transition in the ice plant at the single-cell level using single-cell transcriptomics. This approach provides a mesophyll-specific view of circadian dynamics during CAM induction, a critical survival mechanism under stress. By focusing on transcriptional changes in photosynthetic cells, our research eliminates background noise from non-photosynthetic cells, offering insights into the circadian clock’s association with the 24-hour CAM during the early stages of its induction in the ice plant. The insights gained from this study not only further our understanding of CAM induction, but may also inform the development of stress-tolerant crops.

## RESULTS

### A high-quality ice plant genome sequence and annotation

To enable the single-cell transcriptome analysis of the C_3_-to-CAM transition in the common ice plant, we *de novo* sequenced, assembled and annotated the *M. crystallinum* genome. Long sequences were generated using the Oxford Nanopore Technology PromethION, resulting in reads with an N50 of 12.8kb and coverage of the genome equivalent to 175× (**Supplementary table 1)**. Short DNA reads obtained through Illumina sequencing, achieving a coverage of 120×, were utilized to refine and correct errors in the draft genome assembly. The final assembly had a total length of 368 Mb distributed in 289 contigs, an assembly N50 of 7.19 Mb (n=15), and a BUSCO score of 98.3% using the *embryophyta* dataset as reference, indicating high genome completeness (**Supplementary table 1**). To produce a robust annotation, the ice plant transcriptome was sequenced using PacBio Iso-Seq, generating highly accurate full-length transcripts. The genome annotation process yielded a confident identification of 20,739 genes in the ice plant genome. This gene count is lower than reported for another recently sequenced, chromosome-level assembly of this species^10^. The high degree of similarity between both genome sequences (**Supplementary Figure 1**) confirms the comprehensiveness and accuracy of the sequence we generated. The higher gene count reported previously resulted in large part from the annotation of ribosomal RNAs as protein-coding sequences (see Methods; **Supplementary table 7**), while our incorporation of an IsoSeq dataset significantly improved the precision in identifying protein-encoding genes^15^.

### Single-cell genomics of the ice plant leaf deconstructs the cellular complexity of a facultative CAM organ

A daily evaluation of CAM activity in the leaves of salt-treated and well-watered plants was conducted to define the timing of the switch between C_3_ and CAM under our growing conditions. Because CAM is in part defined by the nocturnal accumulation of malic acid in the vacuoles^4^, we measured differences in titratable leaf acidity between salt-treated and well-watered plants at the beginning of the photoperiod. Our results show that a significant difference in acidity was observed at and after 8 days of the start of the salt treatment (t-test, p-value=5.56×10^−4^; **Supplementary Figure 2**), indicating the onset of a transition from C_3_ to CAM^3,5^. To characterize the transcriptional profile of this transition, we isolated nuclei from 5-week-old ice plant leaves, after 8 days of salt treatment, for single-cell transcriptome analysis. Leaf samples were collected 30 minutes after the onset of the dark phase (Dusk Salt sample) and 30 minutes after the onset of the photoperiod (Dawn Salt sample). In addition, we isolated leaf nuclei from ice plants watered without NaCl at the same time points (Dusk Control and Dawn Control samples). Droplet-based snRNA-seq was performed for all samples using the 10× Genomics platform. Subsequent data processing yielded an integrated, normalized, batch-corrected, and clustered dataset of 17,994 high-quality nuclei grouped into 17 clusters (**Figure 1A and Figure 2B**).

**Figure 1:**
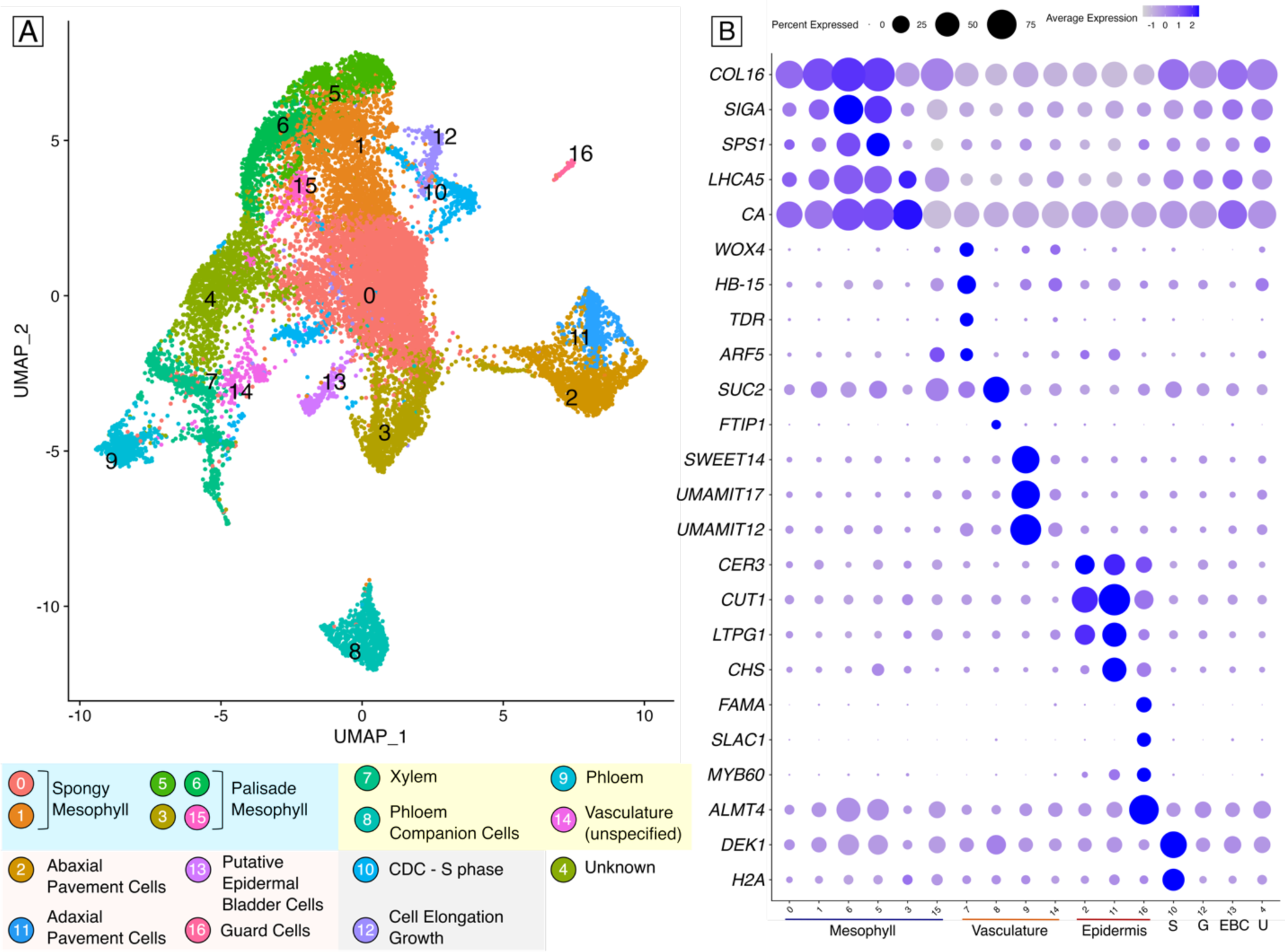
A) UMAP clustering of the integration of all four snRNAseq datasets (Dawn Control and Salt, Dusk Control and Salt). A total of 17,994 nuclei are divided into 17 clusters. B) Dotplot representing the expression of orthologs of cell type-specific markers in the integrated ice plant snRNA-seq dataset. CDC = Cell division cycle, S = S-phase of the cell cycle, G = Growth, EBC = Epidermal Bladder Cells, U = Unknown.

**Figure 2:**
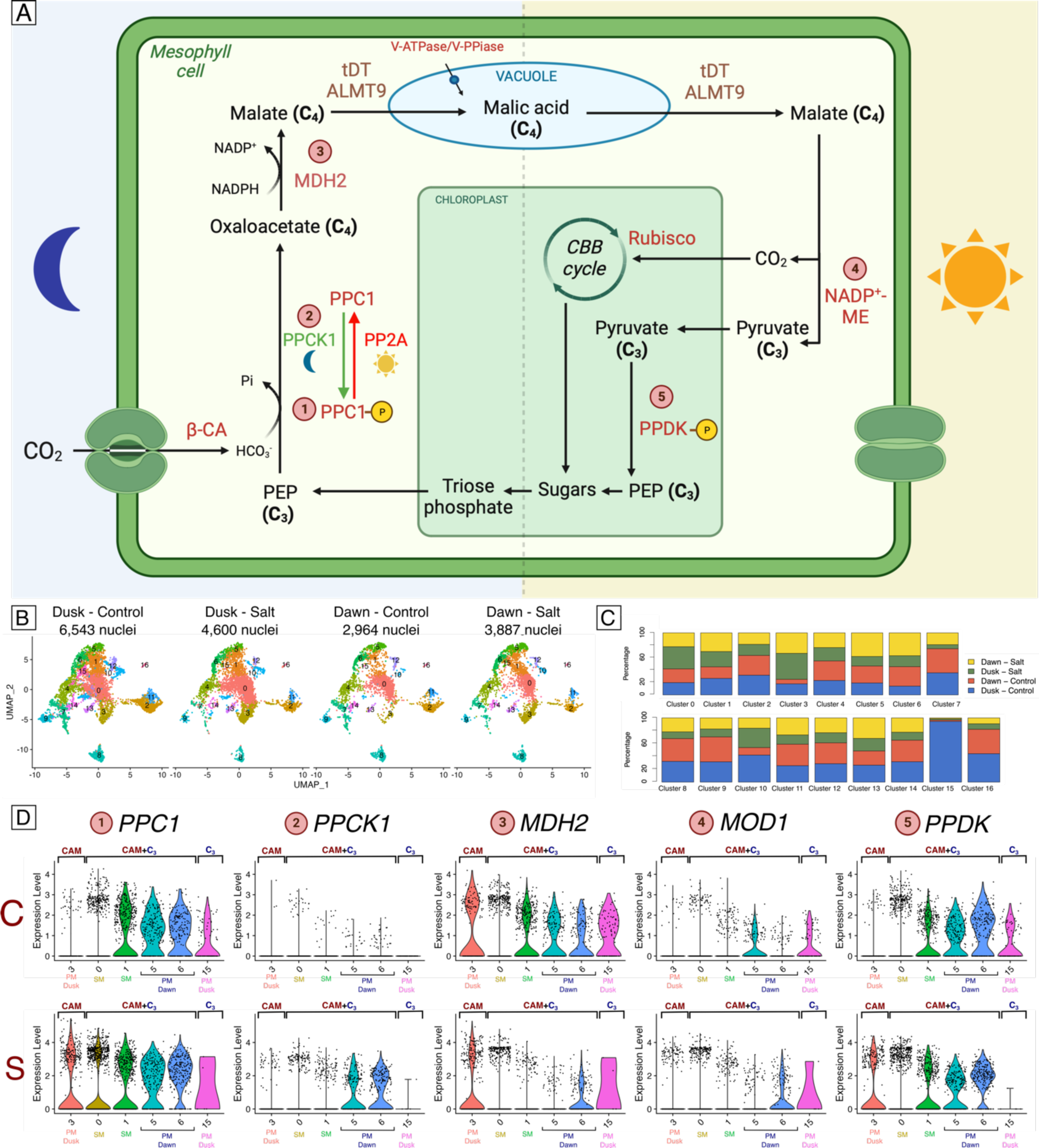
A) Overview of the main biochemical events of the CAM during the 24-hour diel cycle in CAM-performing ice plant mesophyll cells. During the night, atmospheric CO_2_ is converted to HCO_3_^−^ by beta-carbonic anhydrase (β-CA). This product is then added to phospho*eno*lpyruvate (PEP) by PEP carboxylase 1 (PPC1) to generate oxaloacetate (OAA). PPC1 is activated at night via phosphorylation by PPC kinase 1 (PPCK1), and deactivated during the day via dephosphorylation by protein phosphatase 2A (PP2A). OAA is rapidly reduced in the cytosol by malate dehydrogenase (MDH, encoded by *MDH2* in the ice plant), leading to the accumulation of malic acid. Malate is then transported into large vacuoles where it is stored for the remainder of the dark phase. During the daytime, stomata close to minimize water loss by transpiration, and malate is transported out of the vacuole. Subsequently, it is decarboxylated in the cytosol by NAD(P)-malic enzyme (NADP-ME, encoded by *MOD1* in the ice plant), or NAD-malic enzyme, providing CO_2_ for daytime fixation via Rubisco in the Calvin-Benson-Bassham (CBB) cycle. The pyruvate product of decarboxylation is converted to PEP by PPDK. B) UMAP clusterings representing the cells of each sample in the integration C) Sample repartition in the 17 clusters of the integrated dataset. For accurate comparison, the numbers of nuclei per cluster were adjusted based on the total number of nuclei of the sample of origin. D) Violin plots depicting cluster-specific expression profiles of CAM genes in the mesophyll cells of the control samples (C = Dusk Control + Dawn Control, top) and of the salt-treated samples (S = Dusk Salt + Dawn Salt, bottom). The numbers in front of the gene names correspond to the numbers in **Figure 2A**.

Cluster annotation was performed by examining the expression pattern of orthologs of known cell type-specific markers in the ice plant snRNA-seq dataset (**Figure 1B; Supplementary table 2)**. The expression patterns of vasculature-specific markers *WOX4* (Mcr-019817)^16^, *HB-15* (Mcr-005975)^17^, *TDR* (Mcr-013120)^18^, *ARF5* (Mcr-011164)^19^, and *SWEET14* (Mcr-015099)^20^, *UMAMIT12* (Mcr-018247)^21^, *UMAMIT17* (Mcr-001120)^21^ was used to designate clusters 7 and 9 as xylem and phloem cells, respectively. Cluster 8 was found to comprise phloem companion cells (*SUT1* (Mcr-001196)^22^, *FTIP1* (Mcr-015504)^22^), and cluster 14 was annotated as unspecified vasculature cells based on similarity with the expression profiles of both phloem and xylem clusters. The expression of known pavement cell (PC)-specific markers such as *CER3* (Mcr-008261)^23^, *CUT1* (Mcr-001887)^24^, and *LTPG1*(Mcr-004691)^25^ mostly localized to clusters 2 and 11. Refinement of the annotation of these clusters was facilitated by analyzing the expression of the adaxial PC-specific marker, CHS (Mcr-010818)^26^ which exhibited specificity to cluster 11. As a result, cluster 11 was categorized as adaxial PC and cluster 2 as abaxial PC, given the lower expression of CHS. The identification of the guard cell cluster (cluster 16) was based on the specific expression of *FAMA* (Mcr-018185)^27^, *MYB60* (Mcr-019604)^28^, and *SLAC1* (Mcr-006914)^29^. Moreover, the expression of *ALMT4* (Mcr-005476), which was shown to be limited to guard and mesophyll cells^30^, was most pronounced in clusters 16, 5, and 6. The distinct expression of DNA replication markers such as *DEK1* (Mcr-015108)^31^ and *H2A* (Mcr-014584)^32^ in cluster 10 led us to classify it as nuclei within the S-phase of the cell division cycle (CDC).

The abundance of mesophyll cells in leaf tissue, coupled with the diversity of treatments and conditions that impact their transcriptome state, resulted in their assignment to the annotation of multiple clusters. The expression profiles of mesophyll-specific markers *SPS1* (Mcr-010463)^33^, *SIGA* (Mcr-013069)^34^, and *COL16* (Mcr-015233)^35^ identified clusters 0, 1, 5, and 6 as encompassing mesophyll cells. These genes also showed a comparatively higher expression in cluster 4 relative to non-mesophyll cells. However, the low overall signal for each cell-type-specific marker tested in our analyses in cluster 4, combined with the lack of clear transcriptional identity after further investigation of the genes specifically expressed in this cell group, suggested that it contained a mixture of cell types (**Figure 1A and B**). Consequently, cluster 4 could not be classified as a true mesophyll group and was annotated as “unknown”.

Leaves of dicotyledonous species consist of two layers of mesophyll cells. The spongy mesophyll (SM), positioned on the abaxial leaf surface, is composed of cells that display irregular morphological features and serve a crucial function in CO_2_ diffusion within the leaf ^36^. The palisade mesophyll (PM) layer, on the adaxial side, is characterized by thick and tightly packed cells. To distinguish the SM and PM layers, we examined the expression profile of genes involved in light-capture and CO_2_ fixation, which were previously found to be upregulated in the PM^25^. The expression of photosynthesis-associated genes such as *LHCA5* and *CA* facilitated the identification of clusters 3, 5, and 6 as comprising mesophyll cells of the PM layer. Clusters 0 and 1 exhibited a lower level of gene expression associated with light-capturing functions compared to clusters 3, 5, and 6, and the presence of mesophyll-specific markers in these clusters led us to the conclusion that they corresponded to the SM layer. In addition, the relatively elevated expression of certain mesophyll-specific markers in cluster 15, combined with the abundance of differentially expressed (DE) genes encoding chloroplast-localized products strongly indicated its composition of mesophyll cells (**Supplementary table 3**). Further examination of the transcripts defining this cluster highlighted genes known to be prominently expressed exclusively in the evening and nighttime, such as *LEA5* (Mcr-018008)^37^ and *TOC1* (Mcr-012028)^38^. This was also true for genes associated with the *Arabidopsis* evening complex (EC), including *LUX* (Mcr-014497), and *ELF4* (Mcr-002654)^38^ (**Figure 3D**). Pairing these insights with the observation that the vast majority of nuclei in cluster 15 originated from the Dusk – Control sample (**Figure 2C**) supports that cluster 15 represented nighttime mesophyll cells.

**Figure 3:**
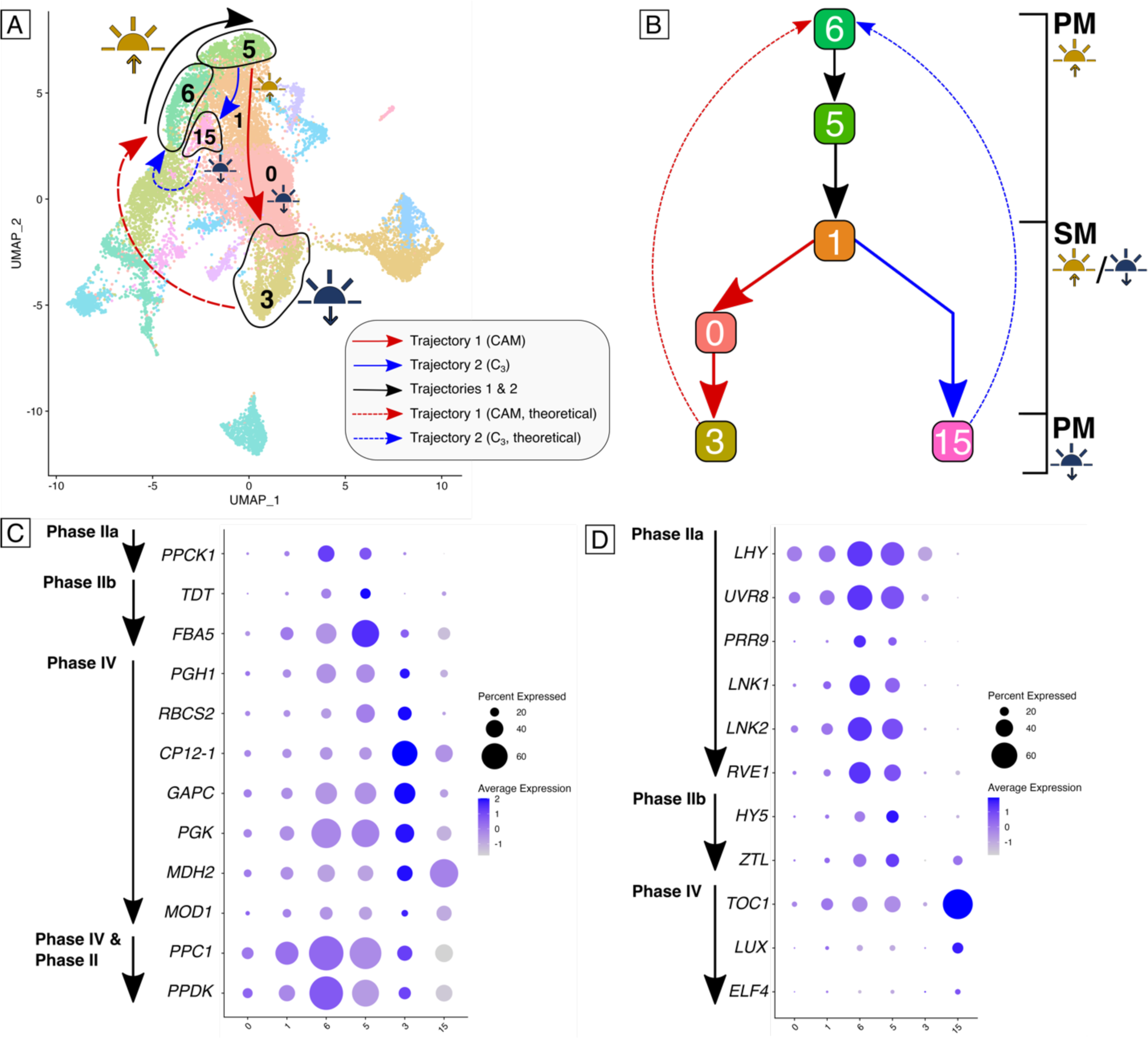
A) Cell trajectories determined by Slingshot on the UMAP clustering of the integrated dataset. The relevant mesophyll cell clusters within the trajectory are annotated by black numerals, indicating their respective cluster numbers. Dotted arrows in our representation symbolize the theoretical return to the transcriptional state characteristic of cells at dawn, a process not explicitly delineated by the trajectories computed using Slingshot. B) Topological representation of the trajectories. C) Cluster-specific expression of a selection of genes involved in the CAM pathway. The phases annotated on the left correspond to the phases as described by Osmond, 1978^12^. D) Cluster-specific expression of a selection of regulators of the plant circadian clock.

Upon annotating the majority of clusters using orthologs of established cell type-specific markers, the identities of clusters 12 and 13 remained ambiguous. For cluster 12, the prevalence of genes associated with auxin metabolism, growth, and cell elongation hinted at cells characterized by robust growth, potentially resembling leaf meristem or G1/G2-phase cells in CDC. However, the use of known G1/G2 and meristematic markers did not identify specific expression within cluster 12 in our analysis, thus precluding a more precise cluster annotation. Given the predominant expression of genes linked to cell elongation and growth (with an average log2 fold change compared to their expression in other clusters exceeding 2, such as *LNG2* (Mcr-019870)^39^, *ARF19* (Mcr-008385)^40^, and *IAA7* (Mcr-011541)^41^; **Supplementary table 3**), cluster 12 was defined as a cell elongation/growth cluster. The same strategy was employed to gain deeper insights into the unique characteristics of cluster 13. This analysis unveiled a predominant presence of transcripts related to ion exchange, homeostasis, transport, and storage functions. Notably, these features aligned with the transcriptional signature of epidermal bladder cells (EBCs). These specialized trichomes play a crucial role in sequestering excessive salts in *M. crystallinum*, enabling the plant to maintain normal function under high salinity conditions, a pivotal process for salt tolerance^42^. Consequently, we designated cluster 13 as a putative EBC cluster.

### Mesophyll cells exhibit high transcriptomic plasticity at the onset of the C_3_-to-CAM switch

The categorization of mesophyll cells into six distinct clusters prompted us to investigate the transcriptional nuances underpinning such heterogeneity. Employing a non-parametric Wilcoxon rank sum test, we compared transcript levels between nuclei from salt-stressed samples (Dawn and Dusk Salt) to those from well-watered samples (Dawn and Dusk Control) within each mesophyll cluster (specifically, clusters 0, 1, 3, 5, 6, and 15; **Supplementary table 4**). Genes identified as DE (adjusted p-value < 0.05) were categorized based on their average log2 fold-change (log2 fold-change > 1.5 = “ highly upregulated”, 1.5 > log2 fold-change > 1 = “upregulated”, 1 > log2 fold-change > 0= “marginally upregulated”). Lastly, DE genes showing a log2 fold-change less than 0 were classified as “downregulated.” Using this scale as a reference, we identified high upregulation of *PPC1*, *MOD1*, and *PPDK* transcripts in nuclei of cluster 3 derived from salt-treated samples, a trend not detected in other mesophyll clusters (**Supplementary table 4**). In addition, an upregulation in *PPC1* and *MOD1* transcripts was observed in clusters 0 and 6 (**Figure 2D, Supplementary table 4**). Earlier studies investigating the diurnal expression pattern of *PPC1* and *MOD1* in CAM- and C_3_-performing ice plant leaves revealed a distinct spike in their expression at dusk in CAM leaves. Contrastingly, the expression of these genes in C_3_ leaves remained low during this time^3,43^. *PPCK1* transcripts, while not elevated in clusters 0 and 3, were highly upregulated in the nuclei of clusters 5 and 6 from the salt-treated samples (**Figure 2D, Supplementary table 4)**. This is in agreement with previous observations that *PPCK1* expression peaked at the end of the dark phase and dawn, only to fall sharply by dusk^3,43^. Furthermore, *MDH2* was downregulated in salt-treated cells of clusters 5 and 6 (**Figure 2D, Supplementary table 4)**. An examination of each cluster’s sample composition (proportional to the total number of nuclei per sample) showed that 41.9% of nuclei in cluster 3 were derived from the Dusk – Salt sample. Similarly, a predominant fraction of nuclei in clusters 5 and 6 were from the Dawn – Salt sample, at 38.1% and 36.9%, respectively (**Figure 2C**). Such patterns, as well as the observations made previously, indicated that cluster 3 was enriched with dusk CAM mesophyll cells (or phase IV CAM cells, following Osmond’s CAM cycle classification^12^), while clusters 5 and 6 predominantly harbored dawn CAM mesophyll cells (or phase II CAM cells^12^). The salt-induced stress did not lead to any variation in CAM enzyme encoding transcripts within cluster 15, reaffirming its annotation as dusk C_3_ mesophyll cells (**Figure 2D, Supplementary table 4)**. The expression patterns of CAM-related genes in cluster 1, following salt treatment, mirrored the profiles observed in the CAM dawn clusters (5 and 6), albeit at a lower magnitude. Similarly, the increase in expression of CAM enzymes that was evident in cluster 3 was also noted, to a lesser degree, in cluster 0 (**Supplementary table 4)**. These findings suggested that cluster 1 likely contained CAM-performing SM cells at dawn, whereas cluster 0 comprised CAM-performing SM cells at dusk. The more modest upregulation of CAM enzyme transcripts in clusters 0 and 1, as compared to clusters 3, 5, and 6, is consistent with the current understanding that SM cells are less engaged in photosynthesis than PM cells^4^.

While our clustering analysis effectively distinguished between C_3_ and CAM mesophyll cells during dusk, no such separation was achieved for C_3_ and CAM cells at dawn, with both cell states apparently contained within clusters 5 and 6. This blending was expected considering that the light phase of the CAM cycle is predominantly characterized by genes from the C_3_ pathway (**Figure 2A**), hampering the segregation of daytime CAM and C_3_ cells based only on transcriptomic profiles.

### Cell trajectory inference analysis reconstructs a CAM 24-hour cycle in mesophyll cells

The separation of mesophyll cells into clusters enriched in nuclei originating from either the Dawn or Dusk samples led us to explore the possibility of reconstructing a 24-hour CAM cycle. Using a trajectory inference (TI) algorithm, we attempted to order the nuclei from each mesophyll cluster along a day/night gradient based on the cell’s phase at the time of nuclei isolation. To refine the analysis and minimize interference from non-mesophyll cells, only clusters 0, 1, 3, 5, 6, and 15 were utilized for the analysis. Given our goal to initiate the trajectory with cells from the earliest light phase, we assessed the expression patterns of *LHY* (Mcr-001290), a pivotal plant circadian clock regulator with peak expression at dawn^44^. Notably, the highest *LHY* expression was found in cluster 6, followed closely by cluster 5 (**Figure 2D**). Therefore, cluster 6 was defined as the starting point of the trajectory. TI algorithm Slingshot^45^ was used to identify the end cluster(s) and delineate trajectories leading to these endpoints. This analysis identified two distinct trajectories. The first traced a path from cluster 6 to 5, advancing to clusters 1, 0, and concluding at cluster 3. Conversely, the second trajectory started at cluster 6, progressed to clusters 5 and 1, and ended at cluster 15 (**Figure 1B and Figure 2A**). Considering that both trajectories concluded at one of two “dusk” clusters, namely cluster 3 and cluster 15, and our prior analysis showed enrichment of CAM mesophyll cells in cluster 3 and C_3_ mesophyll cells in cluster 15, we evaluated the hypothesis that the first trajectory represented a “CAM path” and the second a “C_3_ path”. To achieve this, we examined the expression patterns of genes with known expression profiles over the 24-hour CAM cycle and recognized roles in the pathway. The analysis revealed that *PPCK1*, a gene peaking in expression at dawn, is more highly expressed in cluster 6 (**Figure 3C**). *TDT* (tonoplast dicarboxylate transporter), responsible for malate transport out of the vacuole during the early daytime CAM reaction^46^ (**Figure 2A**), exhibited an inverse expression profile to *PPCK1*, with a marked expression in cluster 5 and lower expression in cluster 6. *FBA5* (Fructose-bisphosphate aldolase 5), integral to glycolysis and gluconeogenesis that take place during the day-phase of CAM, was more highly expressed in cluster 5. *PGH1*, encoding for Enolase – a crucial element of sugar metabolism – was consistently expressed in clusters 5 and 6, but showed preferential expression in cluster 3. This aligned with its established dusk peak expression in CAM ice plant leaves^3^. Similarly, *PGK* (phosphoglycerate kinase) showed notable abundance in dawn clusters but higher expression levels in cluster 3. Genes like *RBCS2* (ribulose bisphosphate carboxylase small subunit 2), *CP12-1* (CBB cycle protein 12-1), and *GAPC* (Glyceraldehyde-3-phosphate dehydrogenase) are pivotal for the CBB cycle, and displayed pronounced expression in cluster 3. This underscored the notion that the C_3_ component of CAM is activated later in the day, persisting into the early parts of dusk (phase IV) as CAM gets underway. Genes encoding for the CAM enzymes MDH2, MOD1, PPC1, and PPDK displayed heightened expression in cluster 3 during dusk, yet maintained considerable expression in many dawn cells (clusters 5 and 6). The observed gene expression patterns traced a clear trajectory through the diel CAM cycle, underlining that the distinct mesophyll clusters in our study mirrored varied phases of the pathway. The successive expression of these genes across clusters 6, 5, and 3 validated that the first trajectory reconstructed an almost complete CAM cycle. As the dawn phase (or CAM phase II) was separated into two distinct clusters, expression of genes with peak expression shortly after dawn (e.g. *LHY*, *UVR8*; **Figure 3C**) was higher in cluster 6, and both trajectories crossed cluster 5 after cluster 6, we hypothesized that cluster 6 encompassed cells at an earlier stage of dawn compared to cluster 5. A differential gene expression analysis between these clusters showed that numerous genes upregulated in cluster 6 (defined by an average log2 fold-change > 0.75 and an adjusted p-value < 0.05) were involved in the light-capture phase of photosynthesis (such as *LHCA4*, *psaK*, *CAB*), along with genes in carotenoid (*CRTISO*^47^) and chlorophyll (*Chlorophyllide a oxygenase*^48^) biosynthesis pathways (**Supplementary table 3)**. Conversely, genes DE in cluster 5 had broader functions in plant metabolism, including lipid biosynthesis (e.g., *FAR1*^49^, *FATB1*^50^), flavonoid biosynthesis (e.g., *PAL*^51^, *FLS*^52^), sugar metabolism and transport (e.g., *ERD6-like transporters*, *FBA5*), and carbon metabolism (*Rubisco small subunit 2*; **Supplementary table 3).** These observations highlighted the temporal sequence from light capture, facilitating the downstream activation of the CBB cycle, eventually leading to sugar and broader plant metabolism. Therefore, we designated the nuclei of cluster 6 as being in phase IIa, and those of cluster 5 as being in phase IIb, which follows shortly after dawn (**Figure 3C & D**).

The observation that cluster 15 mostly consisted of nuclei from Dusk Control (**Figure 1E**), and that trajectory 2 bifurcated to cluster 15 rather than cluster 3, strongly suggested that this trajectory captured the C_3_ diurnal cycle. This insight further confirmed the mixture of C_3_ and CAM cells within clusters 5 and 6, as well as the segregation of dawn and dusk SM cells between clusters 1 and 0, respectively. To further support the conclusion that identified cell trajectories represent a virtual 24-hour cycle, we analyzed the expression patterns of pivotal regulators of the plant circadian clock (**Figure 2D**). Genes crucial for sensing dawn and photoperiod, such as *LHY*, *UVR8*, *PRR9*, *LNK1* and *2*, and *RVE1*, showed pronounced expression in clusters 5 and 6, with a particular enrichment in cluster 6. Interestingly, *HY5* and *ZTL*, which also play a role in mediating light responses, were predominantly expressed in cluster 5. In contrast, evening-specific gene *TOC1* and components of the EC, *LUX* and *ELF4*, exhibited elevated expression in cluster 15. These genes did not present a similarly heightened expression in cluster 3. This aligns with the documented shift in EC component expression seen in *Kalanchoë fedtschenkoi*, a constitutive CAM species^53^. This data further bolsters the categorization of trajectory 2 as representing the “C_3_ path”.

Our TI findings offer an opportunity to juxtapose the temporal shifts in gene expression between CAM and C_3_ diurnal cycles. To provide an additional layer of evidence regarding the distinct nature of each trajectory, we used tradeSeq^54^ to model gene expression changes along both trajectories (**Figure 4**). In addition, a multivariate Wald test was implemented to determine whether the expression levels of CAM-specific genes exhibited significant differences between trajectory 1 and trajectory 2. This analysis validated that CAM genes such as *PPC1*, *MDH*, *PPDK* and *MOD1* were significantly upregulated in trajectory 1 in comparison with trajectory 2 (median fold-change > 0, p-value < 0.05, **Supplementary table 4, Figure 4**).

**Figure 4:**
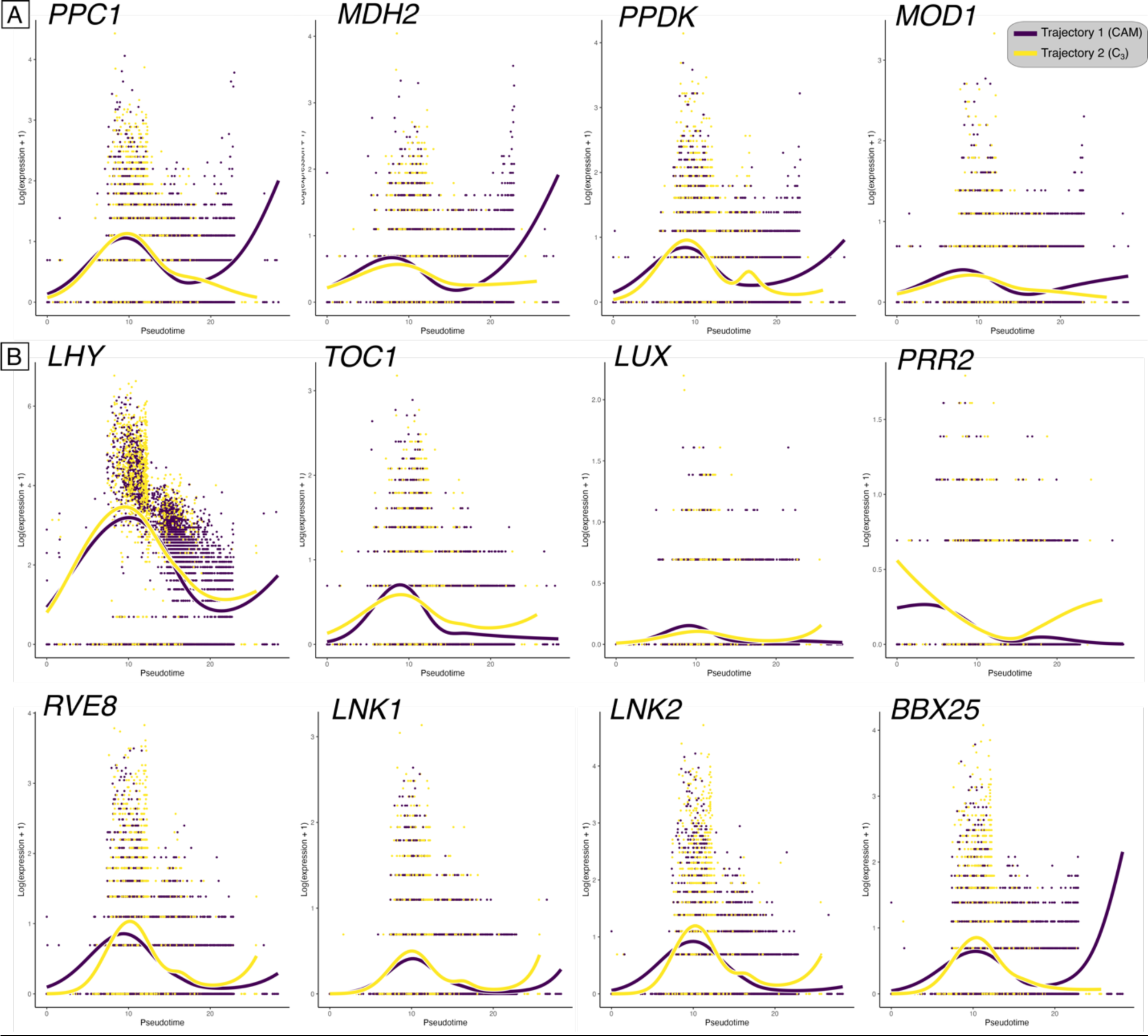
Smoother plots illustrating the expression of select CAM genes (A) and circadian clock genes B) across the two trajectories. Each point on the graph represents a cell within the trajectory, with its position on the y-axis determined by the expression level of the given gene. The x-axis depicts the cell’s advancement along the designated trajectories according to their pseudotime, starting at pseudotime 0 for the onset of the trajectory.

### The CAM and C_3_ mesophyll trajectories are characterized by divergent expression profiles of key circadian clock genes

Examination of pseudotime expression profiles of circadian clock genes along the trajectories underscored important day-night transcript fluctuations. While the expression of *LHY* remained highly similar in both trajectories, other core circadian clock components and genes associated with circadian regulation in plants showed altered expressions in the CAM pathway (**Figure 4B**). The multivariate Wald test indicated that genes such as *TOC1*, *LUX*, *PRR2*, *RVE8*, *LNK1*, and *LNK2* were differentially expressed between the C_3_ and CAM trajectories (median fold-change > 0, p-value < 0.05; **Supplementary table 4, Figure 4B**). This observation was consistent with results from an earlier study that compared the expression profiles of several of these genes across a 24-hour period between the obligate CAM species *K. fedtschenkoi* and the C_3_ species *A. thaliana*^53^. Such findings validated that our trajectories accurately reflected the 24-hour cycles of CAM and C_3_, and also suggested that reduced expression of key circadian genes was linked to the CAM phenotype. Interestingly, *BBX25*, a transcription factor known to inhibit photomorphogenesis in Arabidopsis by interacting with *HY5* and implicated in various plant circadian regulation processes^55^, showed significant upregulation at the end of the CAM trajectory but remained low in the C_3_ trajectory (**Figure 4B**, median fold-change = 0.87, waldStat=21.26, p-value=9.28E-05). Although our study did not definitively confirm the hypothesis that *BBX25* is involved in the downregulation of other circadian clock genes, continued and more detailed research into the function of *BBX25* across different CAM species could potentially uncover its role in the CAM induction process.

## DISCUSSION

Research on CAM has been impeded by an overall lack of genetic and genomic resources. Major questions, such as the evolutionary origin of the pathway, remain to be answered^56^. Comparative genomic analyses represent a powerful approach for addressing this matter, but require access to abundant genomic data. In this regard, our publicly available high-quality annotation of the *M. crystallinum* genome constitutes a valuable new resource for the elucidation of CAM origins. To confirm the quality of the genome generated in this study, a comparative analysis was conducted with the only previously published *M. crystallinum* genome (referred to as IcePlant1 for clarity). This analysis revealed discrepancy in the number of protein-coding genes between genomes: 20,739 in our genome (referred to as IcePlant2) versus 24,234 in IcePlant1, largely due to misannotation of ribosomal RNA (rRNA) sequences as protein-coding genes in IcePlant1 (see **Methods**, **Supplementary table 7)**. This observation, if confirmed, would suggest that the higher count of protein-coding genes reported in IcePlant1 may, in part, stem from a misannotation of these rRNA sequences. Consequently, the methodologies employed in IcePlant2, such as full-length transcript sequencing with IsoSeq, appear to have provided a more accurate representation of the ice plant’s genomic structure.

The first single-cell transcriptome atlas of a CAM species provides a basis for elucidating the underlying molecular mechanisms and processes behind cellular function and diversity in an organism capable of surviving under extreme salinity and drought conditions. Despite the inherent recalcitrance of the ice plant and other CAM species to genetic transformation^58^, hindering the direct validation of cell cluster annotations, the employment of orthologs to well-established cell type-specific markers has resulted in the accurate annotation of cells across different plant species^22,25,27,34,59^. Furthermore, the consistent and highly specific expression patterns of these markers within our identified cell clusters further reinforced the reliability of their annotations (**Figure 1B**). Our findings indicate that only SM and PM mesophyll cells display a significant response to salt stress in inducing CAM under our experimental conditions. Epidermal and vascular cell-types may undergo less dramatic transcriptional changes or present a delayed response to the stimulus. This implies that future efforts to engineer CAM traits into C_3_ species should be targeted at mesophyll cells. Such a targeted approach could minimize pleiotropic effects and prevent the disruption of crucial cellular processes in other cell types. Repeating a similar analysis using nuclei extracted from leaves in a more advanced phase of transition could further distinguish the transcriptional profile of CAM cells, making it less likely for C_3_ and CAM cells to cluster together. Yet, capturing the transcriptome from cells at the early stages of the transition may enable the discovery of regulators of CAM induction. Such insights could pave the way for significant strides in the effort to introduce CAM traits into drought-sensitive crops.

Our research has uncovered distinct expression patterns for several key components and regulators of the circadian clock between the two distinct mesophyll cell trajectories (**Figure 5**). Core circadian clock genes known to show high transcript accumulation in the evening in the C_3_ plant *Arabidopsis* (e.g., *TOC1* and *LUX*) exhibited noticeable downregulation at dusk specifically in the CAM trajectory (**Figure 4B, Figure 5, Supplementary table 4**). These expression patterns, though not typical, aligned with findings from a study on the timing of expression of these genes in *K. fedtschenkoi* and *Arabidopsis*. This suggests that the downregulation of circadian clock genes plays a central role in the CAM pathway. Although this study did not define the cause for the low expression of circadian clock genes in CAM ice plant leaves, one hypothesis is that these genes are regulated differently in CAM, when compared to C_3_ plants. Thus, the pronounced increase in *BBX25* expression observed in the CAM trajectory calls for additional research to examine its possible involvement in the CAM induction process in the ice plant. Recent evidence showing that overexpressing *Ginkgo BBX25* enhances salt tolerance in *Populus*^60^ suggests that this upregulation might relate to salt stress application. However, it is not to be excluded that the genes involved in salt-stress and drought response could also be crucial in regulating CAM induction in facultative CAM species. The widespread downregulation of circadian clock genes in CAM plants could have significant biological implications, potentially reflecting a necessity to delay or modify the transition to the dark phase compared to C_3_ plants. One hypothesis may be that the expression of certain clock genes which are active in the evening in C_3_ species might contribute to stomatal closure. Because the stomata of CAM plants open at night, it is conceivable that the suppression of these genes may be required to facilitate this unique stomatal behavior.

**Figure 5:**
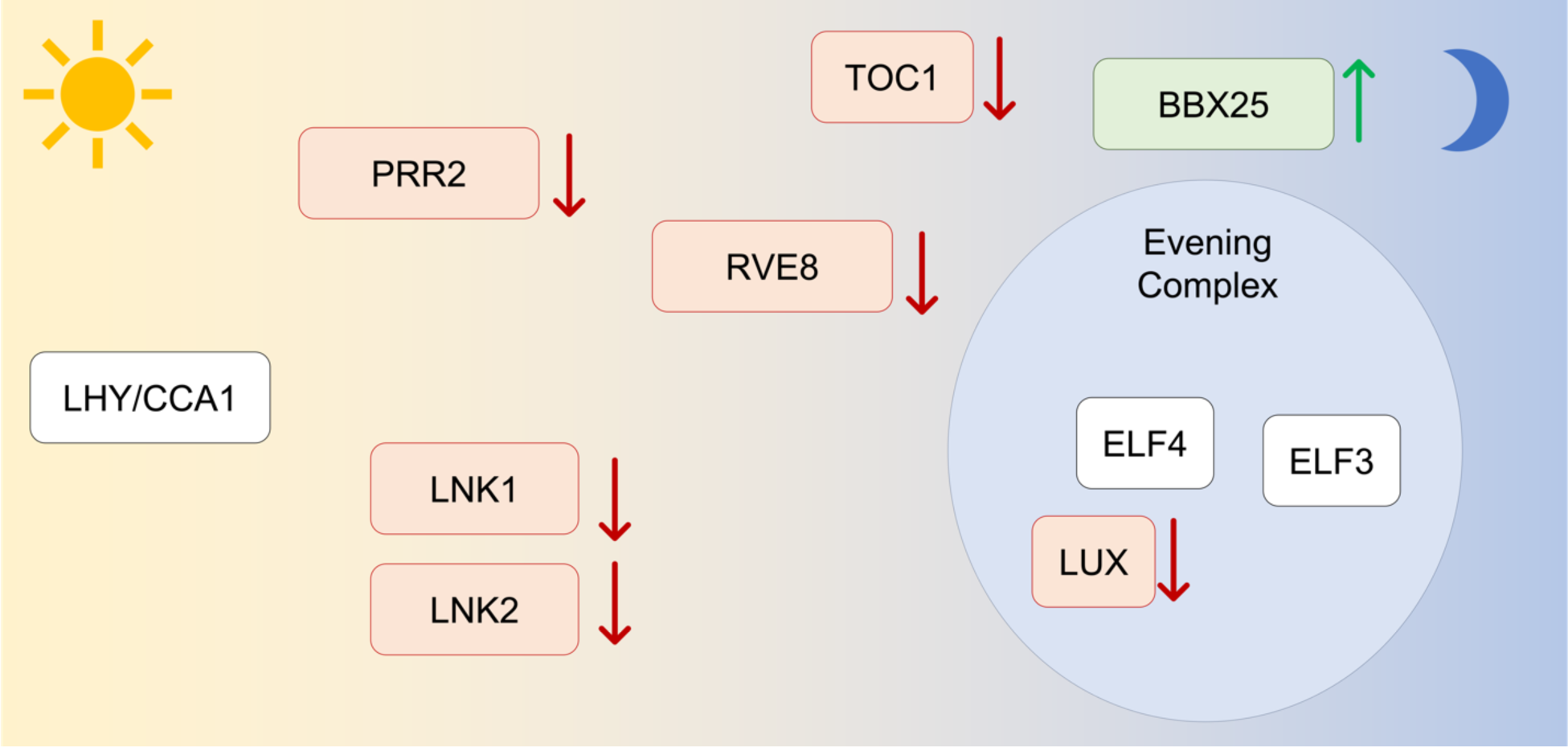
Representation of differentially expressed circadian genes in the CAM trajectory as compared to the C_3_ trajectory. Genes highlighted in red displayed significant downregulation (p-value<0.05, **Supplementary table 4**) in the CAM trajectory, whereas the gene highlighted in green displayed significant upregulation in CAM. Genes with a white background represent those without notable differences in expression between the two trajectories. The diagram uses a left-to-right gradient to symbolize the progression from dawn to dusk. The spatial positioning of each gene within the figure approximates the timing of their expression over the course of the day.

The separation of dawn mesophyll cells into two distinct clusters, specifically clusters 5 and 6, offers an intriguing perspective through which to view the traditionally described four-phase CAM model. Our data suggests a nuanced view of Phase II, typically characterized as the bridge between the nocturnal and diurnal phases. Results from the TI analysis, paired with the expression profiles of crucial light-responsive circadian genes, suggest that the nuclei within cluster 6 are in an earlier state relative to those in cluster 5 (**Figure 3A-D**). Thus, the elevated expression of *MDH2* and *MOD1* in cluster 6 upon salt treatment, as depicted in **Figure 2D**, may be indicative that cluster 6 contains cells on the edge of exiting the nocturnal phase, when the transcript levels for these genes are still abundant, while cluster 5 could exclusively consist of light-acclimated cells. This interpretation is further cemented by the trajectory illustration in **Figure 4**. Even the earliest cells in the trajectory, despite originating from cluster 6, exhibit transcriptomic profiles characteristic of nighttime for the genes displayed. Our findings thus amplify the perspective that the four-phase CAM model may be more plastic than originally proposed^13^. Our data sheds light on the existence of distinct transcriptomic states between phase I and phase III, tentatively labeled as phase IIa and phase IIb in this study.

## CONCLUSION

This research offers new insight into the cellular and molecular dynamics driving CAM induction and adaptation in the ice plant. Notably, we discovered that only mesophyll cells display significant changes in expression at the onset of this process in ice plant leaves. By analyzing transcriptional variations at the single-cell level and computationally reconstructing the C_3_ and CAM diel cycles in mesophyll cells, we found that the CAM induction process significantly influences circadian gene expression. Specifically, we discovered that the transition to CAM in the mesophyll cells of ice plant leaves is characterized by the widespread downregulation of key circadian clock genes at night. In parallel, CAM mesophyll cells express other circadian clock genes in the evening which were not observed in C_3_ mesophyll cells. Delving deeper into these alterations can not only further our understanding of plant stress responses but also present substantial opportunities for agricultural advancements, particularly in enhancing drought and salinity tolerance.

## MATERIALS AND METHODS

### Genome sequencing

High molecular weight genomic DNA was extracted using 40g of fresh tissue following a protocol consisting of nuclei isolation^61^ followed by SDS based isolation^62^ and NaCl cleanup^63^. Long-reads were sequenced using the Oxford Nanopore Technology (UK) PromethION platform, and the library was constructed according to the respective manufacturer’s protocols. The Covaris g-TUBE was used to shear genomic DNA into fragments between 6 and 20 kb. Fragments from the nanopore library were the size selected at 12 kb using Electrophoretic Lateral Fractionator (ELF). The resulting coverage was 175X and the N50 was 12.8kb. Short-reads were sequenced with the Illumina (USA) NovaSeq 6000 platform, and the library was constructed according to the respective manufacturer’s protocols. The coverage for the short Illumina reads was 120X.

### Genome assembly, polishing, and annotation

The output reads from the PromethION were assembled using FLYE^64^, an assembler designed for nanopore reads, using a reduced coverage of 50X by selecting the longest fragments. The resulting assembly was polished using Pilon^65^, achieving a genome length of 368.0 Mb divided into 289 contigs, and a contig N50 of 7.19 Mb (n=15). A Benchmarking Universal Single-Copy Orthologs (BUSCO)^66^ score using the *embryophyta* dataset was calculated to evaluate genome completeness. *de novo* genome annotation was carried out using Maker2^67^. Only models with an Annotation Edit Distance (AED) < 1 and one or more PFAM domains were considered.

### Transcriptome sequencing

The transcriptomic profile of the ice plant was generated utilizing PacBio Iso-seq. RNA was extracted from 57 plants at diverse developmental stages, ranging from seedling to flowering, grown under two distinct conditions (well-watered and subjected to salt stress), during both diurnal and nocturnal periods. This extraction included leaves, stems, and roots, and followed the extraction protocol established by Chang et al., 1993^68^. The subsequent IsoSeq analysis was conducted on the PacBio SEQUEL IIe platform at the Interdisciplinary Center for Biotechnology Research (ICBR) at the University of Florida. The cDNA was size selected and put into two groups: those ≤ 2.5 kb and those > 2.5 kb in length. Fragments larger than 2.5 kb were subjected to reamplification, and subsequently, cDNA from both size categories were pooled equimolarly in preparation for sequencing. A single SMRT (Single-molecule real-time sequencing) cell run was performed with a movie time of 30 hours.

### Genome comparison

Genome-wide alignments for genome comparison and dot plots were generated through the D-GENIES web application^57^ using the Minimap2^69^ v2.24 software package. Analysis of the genes present on the contigs unique to IcePlant1 was performed using BLASTN analysis against the nucleotide NCBI database with an e-value threshold of 1×10^−05^. Alignment of the whole genomes highlighted the presence of unique contigs in IcePlant1 (**Supplementary table 7**). A closer examination of these contigs, specifically regarding the genes they harbor, revealed that many of these genes, annotated as protein-coding in IcePlant1, have significant matches to rRNAs (**Supplementary table 7)**. Numerous instances of these rRNA-encoding sequences were identified across IcePlant1, some presenting over a hundredfold replication of the same coding DNA sequence, all annotated as unique protein-coding genes. Moreover, BLAST searches using the same queries found targets in the IcePlant2 assembly, but not in the IcePlant2 CDS (coding sequence) file. This confirms the absence of rRNA sequences in the genes annotated as protein-coding in IcePlant2.

### Plant material growth conditions

Seeds of *M. crystallinum* were germinated in moistened soil, and seedlings were transferred to 946mL containers one week after germination. Plants were placed in a growth chamber under 150 μmol m^−2^ s^−1^ white lights to avoid CAM induction due to high photosynthetic photon flux density. The chamber had a 12h/12h light/dark cycle with temperatures of 26°C during the light phase and 18°C during the dark phase. After transplantation, plants were watered daily with 50mL of 0.5X Hoagland’s solution. Four weeks after sowing, a group of plants was watered daily with 50mL 0.5X Hoagland’s solution containing 0.5M NaCl to induce CAM.

### Titratable acidity

Leaf nocturnal acidification was measured in both well-watered and salt-treated plants following a previously described method ^3^. Leaves were collected at the start of the light period (hour 0), and 2g of fresh mass per sample were homogenized in 80% methanol. The leaf tissue homogenate was then titrated against 5 mM NaOH until reaching pH 7 using a pH meter. For each condition, leaves were collected from three biological replicates at each time-point for a total of 6 biological samples per time-point.

### Sample collection for snRNA-seq

For snRNA-seq analysis, the third pair of leaves was collected from 36-day old plants that were either well-watered or salt-treated for 8 days. Samples were collected during both the light and dark phases, resulting in four samples at 36 days. The same collection procedure was applied to the unique 28-day old sample (Day 0 of salt-treatment). Tissue collection was performed in a walk-in cold chamber at 4°C, followed by nuclei isolation.

### Nuclei isolation from *M. crystallinum* leaves for snRNA-seq

Nuclei isolation was performed following a previously described method^70^. All buffers and materials used were pre-cooled at 4°C, and the entire procedure was conducted at the same temperature. For the dark-phase samples, the procedure was performed under darkness, with only an overhead green light used as the light source. Entire leaves were fragmented on a glass plate in 200 μL of Nuclei Isolation Buffer^70^ containing 0.5 U/mL Protector RNase Inhibitor (Sigma Aldrich) using a razor blade. Nuclei in solution were then transferred to a 50 mL tubed containing NIB and placed onto a rotating shaker for 5 minutes. Homogenates were then successively filtered through miracloth (Calbiochem), and then 40 μm and 20 μm filters. Samples were centrifuged at 600g for 5 minutes and washed two times using NIB WASH buffer^70^. The resulting pellets were resuspended in 450 μL NIB WASH, and filtered one last time using 40 μm filters.

### Nuclei sorting

Following nuclei isolation, the samples were stained with 5 μg/mL DAPI and incubated for 5 minutes at room temperature. The nuclei were then isolated from the cell debris in suspension using Fluorescence Activated Nuclei Sorting (FANS) utilizing the BD FACSAria™ IIU/III at ICBR at the University of Florida. The nuclei sorting settings employed in this study have been previously described^70^.

### cDNA synthesis and library preparation

Gel Beads in Emulsion (GEM) generation, cDNA synthesis and library construction were carried out as indicated by the 10X Genomics Chromium Next GEM Single Cell 3ʹ Reagent Kits v3.1 user guide. For each sample, 10X microfluidic chips were loaded with 10 thousand nuclei, and 13 PCR cycles were used for cDNA amplification. Subsequent cDNA library sequencing was carried out on the NovaSeq 6000 System at the ICBR, with S1 flow cell 2×100 sequencing kit but with cycling of 28 read 1, 10 index 1, 10 index 2, and 90 read 2. For the Dusk Salt sample, a second technical replicate was obtained by re-sequencing of the library a second time to obtain additional reads and equilibrate the total number of nuclei identified in each sample. Both sequencing files from the same Dusk Salt library were concatenated prior to downstream analyses.

### Data processing and clustering

The cDNA sequences were processed and the counts matrix was generated using Cell Ranger (v7.0, 10× Genomics) with default parameters. For each sample, an expected number of 10 thousand cells were considered using the option “--expect-cells”. The raw data from Cell Ranger was used for subsequent quality control and filtering steps. First, we took measures to eliminate potential contamination from chloroplast cDNA. To achieve this, we removed genes with high similarity (BLASTN, e-value < 1×10^−8^) to the ones encoded by the *M. crystallinum* chloroplast genome (NCBI NC_029049) from the raw counts matrix (**Supplementary table 8**). To identify empty droplets and low-quality cells, we used the emptyDrops function from the DropletUtils package (version 1.20.0). We ensured the appropriateness of the model for removing empty droplets by testing it on each sample, as per the recommendations in the package documentation, using the “test.ambient=TRUE” option. Under the null hypothesis, p-values for empty cells (cells with total counts below the set threshold) should distribute uniformly. Therefore, we applied various UMI count thresholds to each sample until we achieved a uniform distribution of p-values for cells with low total counts. This approach allowed us to determine a unique minimum UMI threshold for each sample, effectively removing low-quality nuclei while preserving biologically relevant nuclei. Details of the thresholds applied and the nuclei retained can be found in **Supplementary table 9**. Any gene that was not detected in at least three cells was excluded from the Seurat objects. Data pre-processing and analysis steps, such as normalization, and batch-effect removal, were conducted using Seurat V4.3.0. Integration and clustering of the four snRNAseq datasets was carried out at a resolution of 0.55. Uniform Manifold Approximation and Projection (UMAP) was used to reduce the complexity and visualize the data in two dimensions. 30 components were consistently selected for dimensionality reduction across all samples.

### Cluster annotation

Cell-type identification in our datasets was performed by examining the expression pattern of orthologs in the ice plant genome of known cell-type specific markers from the literature (**Supplementary table 2**). For improved cluster annotation, we used Seurat to identify the top 50 genes with the most specific expression in each cluster.

### Trajectory inference analysis

Nuclei from mesophyll clusters 0, 1, 3, 5, 6, and 15 were ordered along pseudotime using Slingshot version 2.6.0^45^. TradeSeq version 1.12.0 was employed to generate gene expression curves along the trajectories and carry out trajectory-based differential expression analyses.

## DATA AVAILABILITY

Raw and processed genomic, IsoSeq, and single-cell transcriptomic ice plant data generated during the study are available on National Center for Biotechnology Information (NCBI).

## ACKNOWLEDGMENTS

The authors extend their heartfelt gratitude to Dr. Klaus Winter from the Smithsonian Tropical Research Institute for his invaluable contributions to this manuscript. His expert insights on CAM significantly enhanced the quality of our work. The authors also thank Mariza Miranda from the Flow Cytometry and Confocal Microscopy core at the Interdisciplinary Center for Biotechnology Research (University of Florida). Finally, the authors thank Dr. Sixue Chen from the University of Mississippi Department of Biology for providing *M. crystallinum* seeds.

## AUTHOR CONTRIBUTIONS

Conceptualization, NP and MK; Methodology, NP, CD, WJP, and MK; Software, NP; Formal Analysis, NP; Investigation, NP; Data Curation, NP; Visualization, NP; Writing – Original Draft, NP; Writing – Review & Editing, NP, WJP and MK; Resources, MK and BB; Supervision, CD and MK; Project Administration, NP, CD, and MK.

## REFERENCES

1. Gilman, I. S. et al. The CAM lineages of planet Earth. Annals of Botany mcad135 (2023) doi:10.1093/aob/mcad135.

2. Edwards, E. J. Evolutionary trajectories, accessibility and other metaphors: the case of C4 and CAM photosynthesis. New Phytologist 223, 1742–1755 (2019).

3. Cushman, J. C., Tillett, R. L., Wood, J. A., Branco, J. M. & Schlauch, K. A. Large-scale mRNA expression profiling in the common ice plant, Mesembryanthemum crystallinum, performing C3 photosynthesis and Crassulacean acid metabolism (CAM). Journal of Experimental Botany 59, 1875–1894 (2008).

4. Borland, A. M. et al. Engineering crassulacean acid metabolism to improve water-use efficiency. Trends in Plant Science 19, 327–338 (2014).

5. Winter, K. & Smith, J. A. C. CAM photosynthesis: the acid test. New Phytologist 233, 599– 609 (2022).

6. Borland, A. M., Griffiths, H., Hartwell, J. & Smith, J. A. C. Exploiting the potential of plants with crassulacean acid metabolism for bioenergy production on marginal lands. Journal of Experimental Botany 60, 2879–2896 (2009).

7. Dodd, A. N., Griffiths, H., Taybi, T., Cushman, J. C. & Borland, A. M. Integrating diel starch metabolism with the circadian and environmental regulation of Crassulacean acid metabolism in Mesembryanthemum crystallinum. Planta 216, 789–797 (2003).

8. Winter, K. & von Willert, D. J. NaCl-induzierter crassulaceensäurestoffwechsel bei Mesembryanthemum crystallinum. Zeitschrift für Pflanzenphysiologie 67, 166–170 (1972).

9. Winter, K. & Holtum, J. A. M. Facultative crassulacean acid metabolism (CAM) plants: powerful tools for unravelling the functional elements of CAM photosynthesis. Journal of Experimental Botany 65, 3425–3441 (2014).

10. Shen, S. et al. High-quality ice plant reference genome analysis provides insights into genome evolution and allows exploration of genes involved in the transition from C3 to CAM pathways. Plant Biotechnol J (2022) doi:10.1111/pbi.13892.

11. Perron, N., Tan, B., Dufresne, C. P. & Chen, S. Chapter Thirteen - Proteomics and phosphoproteomics of C3 to CAM transition in the common ice plant. in Methods in Enzymology (ed. Jez, J.) vol. 676 347–368 (Academic Press, 2022).

12. Osmond, C. B. Crassulacean Acid Metabolism: A Curiosity in Context. Annual Review of Plant Physiology 29, 379–414 (1978).

13. Osmond, B., Neales, T. & Stange, G. Curiosity and context revisited: crassulacean acid metabolism in the Anthropocene. Journal of Experimental Botany 59, 1489–1502 (2008).

14. Nohales, M. A. Spatial Organization and Coordination of the Plant Circadian System. Genes (Basel) 12, 442 (2021).

15. Hu, T., Chitnis, N., Monos, D. & Dinh, A. Next-generation sequencing technologies: An overview. Human Immunology 82, 801–811 (2021).

16. Ji, J. et al. WOX4 Promotes Procambial Development. Plant Physiology 152, 1346–1356 (2010).

17. Ohashi-Ito, K. & Fukuda, H. HD-Zip III Homeobox Genes that Include a Novel Member, ZeHB-13 (Zinnia)/ATHB-15 (Arabidopsis), are Involved in Procambium and Xylem Cell Differentiation. Plant and Cell Physiology 44, 1350–1358 (2003).

18. Fisher, K. & Turner, S. PXY, a Receptor-like Kinase Essential for Maintaining Polarity during Plant Vascular-Tissue Development. Current Biology 17, 1061–1066 (2007).

19. Konishi, M., Donner, T. J., Scarpella, E. & Yanagisawa, S. MONOPTEROS directly activates the auxin-inducible promoter of the Dof5.8 transcription factor gene in Arabidopsis thaliana leaf provascular cells. Journal of Experimental Botany 66, 283–291 (2015).

20. Fei, H. et al. OsSWEET14 cooperates with OsSWEET11 to contribute to grain filling in rice. Plant Science 306, 110851 (2021).

21. Zhao, C. et al. Detailed characterization of the UMAMIT proteins provides insight into their evolution, amino acid transport properties, and role in the plant. J Exp Bot 72, 6400–6417 (2021).

22. Kim, J.-Y. et al. Distinct identities of leaf phloem cells revealed by single cell transcriptomics. The Plant Cell 33, 511–530 (2021).

23. Kim, H., Yu, S., Jung, S. H., Lee, B. & Suh, M. C. The F-Box Protein SAGL1 and ECERIFERUM3 Regulate Cuticular Wax Biosynthesis in Response to Changes in Humidity in Arabidopsis. The Plant Cell 31, 2223–2240 (2019).

24. Hooker, T. S., Millar, A. A. & Kunst, L. Significance of the Expression of the CER6 Condensing Enzyme for Cuticular Wax Production in Arabidopsis. Plant Physiology 129, 1568–1580 (2002).

25. Tenorio Berrío, R., et al. Single-cell transcriptomics sheds light on the identity and metabolism of developing leaf cells. Plant Physiology 188, 898–918 (2022).

26. Mahroug, S., Courdavault, V., Thiersault, M., St-Pierre, B. & Burlat, V. Epidermis is a pivotal site of at least four secondary metabolic pathways in Catharanthus roseus aerial organs. Planta 223, 1191–1200 (2006).

27. Liu, Z. et al. Global Dynamic Molecular Profiling of Stomatal Lineage Cell Development by Single-Cell RNA Sequencing. Molecular Plant 13, 1178–1193 (2020).

28. Rusconi, F. et al. The Arabidopsis thaliana MYB60 promoter provides a tool for the spatio-temporal control of gene expression in stomatal guard cells. Journal of Experimental Botany 64, 3361–3371 (2013).

29. Vahisalu, T. et al. SLAC1 is required for plant guard cell S-type anion channel function in stomatal signalling. Nature 452, 487–491 (2008).

30. Eisenach, C. et al. ABA-Induced Stomatal Closure Involves ALMT4, a Phosphorylation-Dependent Vacuolar Anion Channel of Arabidopsis. The Plant Cell 29, 2552–2569 (2017).

31. Roeder, A. H. K., Cunha, A., Ohno, C. K. & Meyerowitz, E. M. Cell cycle regulates cell type in the Arabidopsis sepal. Development 139, 4416–4427 (2012).

32. Joo, H.-Y. et al. Regulation of cell cycle progression and gene expression by H2A deubiquitination. Nature 449, 1068–1072 (2007).

33. Chávez-Bárcenas, A. T. et al. Tissue-Specific and Developmental Pattern of Expression of the Rice sps1 Gene1. Plant Physiology 124, 641–654 (2000).

34. Wang, Y., Huan, Q., Li, K. & Qian, W. Single-cell transcriptome atlas of the leaf and root of rice seedlings. Journal of Genetics and Genomics 48, 881–898 (2021).

35. Ohmiya, A., Oda-Yamamizo, C. & Kishimoto, S. Overexpression of CONSTANS-like 16 enhances chlorophyll accumulation in petunia corollas. Plant Science 280, 90–96 (2019).

36. Borsuk, A. M., Roddy, A. B., Théroux-Rancourt, G. & Brodersen, C. R. Structural organization of the spongy mesophyll. The New Phytologist 234, 946 (2022).

37. Karpinska, B. et al. Late Embryogenesis Abundant (LEA)5 Regulates Translation in Mitochondria and Chloroplasts to Enhance Growth and Stress Tolerance. Frontiers in Plant Science 13, (2022).

38. Huang, H. & Nusinow, D. A. Into the Evening: Complex Interactions in the Arabidopsis Circadian Clock. Trends in Genetics 32, 674–686 (2016).

39. Lee, Y. K. et al. LONGIFOLIA1 and LONGIFOLIA2, two homologous genes,regulate longitudinal cell elongation in Arabidopsis. Development 133, 4305–4314 (2006).

40. Lee, K., Park, O.-S. & Seo, P. J. Arabidopsis ATXR2 deposits H3K36me3 at the promoters of LBD genes to facilitate cellular dedifferentiation. Sci Signal 10, eaan0316 (2017).

41. Liscum, E. & Reed, J. W. Genetics of Aux/IAA and ARF action in plant growth and development. Plant Mol Biol 49, 387–400 (2002).

42. Barkla, B. J., Vera-Estrella, R. & Raymond, C. Single-cell-type quantitative proteomic and ionomic analysis of epidermal bladder cells from the halophyte model plant Mesembryanthemum crystallinum to identify salt-responsive proteins. BMC Plant Biol 16, 110 (2016).

43. Lim, S. D., Lee, S., Choi, W.-G., Yim, W. C. & Cushman, J. C. Laying the Foundation for Crassulacean Acid Metabolism (CAM) Biodesign: Expression of the C4 Metabolism Cycle Genes of CAM in Arabidopsis. Frontiers in Plant Science 10, (2019).

44. Lu, S. X., Knowles, S. M., Andronis, C., Ong, M. S. & Tobin, E. M. CIRCADIAN CLOCK ASSOCIATED1 and LATE ELONGATED HYPOCOTYL Function Synergistically in the Circadian Clock of Arabidopsis. Plant Physiol 150, 834–843 (2009).

45. Street, K. et al. Slingshot: cell lineage and pseudotime inference for single-cell transcriptomics. BMC Genomics 19, 477 (2018).

46. Zhang, J. et al. Light-responsive expression atlas reveals the effects of light quality and intensity in Kalanchoë fedtschenkoi, a plant with crassulacean acid metabolism. GigaScience 9, giaa018 (2020).

47. Isaacson, T., Ohad, I., Beyer, P. & Hirschberg, J. Analysis in Vitro of the Enzyme CRTISO Establishes a Poly-cis-Carotenoid Biosynthesis Pathway in Plants. Plant Physiology 136, 4246–4255 (2004).

48. Reinbothe, C. et al. A role for chlorophyllide a oxygenase in the regulated import and stabilization of light-harvesting chlorophyll a/b proteins. Proceedings of the National Academy of Sciences 103, 4777–4782 (2006).

49. Jones, A., Davies, H. M. & Voelker, T. A. Palmitoyl-acyl carrier protein (ACP) thioesterase and the evolutionary origin of plant acyl-ACP thioesterases. Plant Cell 7, 359–371 (1995).

50. Domergue, F. et al. Three Arabidopsis Fatty Acyl-Coenzyme A Reductases, FAR1, FAR4, and FAR5, Generate Primary Fatty Alcohols Associated with Suberin Deposition1[C][W][OA]. Plant Physiol 153, 1539–1554 (2010).

51. González-Mendoza, D. et al. Changes in the phenylalanine ammonia lyase activity, total phenolic compounds, and flavonoids in Prosopis glandulosa treated with cadmium and copper. An. Acad. Bras. Ciênc. 90, 1465–1472 (2018).

52. Ma, G. et al. Molecular characterization of a flavanone 3-hydroxylase gene from citrus fruit reveals its crucial roles in anthocyanin accumulation. BMC Plant Biology 23, 233 (2023).

53. Moseley, R. C. et al. Conservation and Diversification of Circadian Rhythmicity Between a Model Crassulacean Acid Metabolism Plant Kalanchoë fedtschenkoi and a Model C3 Photosynthesis Plant Arabidopsis thaliana. Frontiers in Plant Science 9, (2018).

54. Van den Berge, K., et al. Trajectory-based differential expression analysis for single-cell sequencing data. Nat Commun 11, 1201 (2020).

55. Gangappa, S. N. et al. The Arabidopsis B-BOX Protein BBX25 Interacts with HY5, Negatively Regulating BBX22 Expression to Suppress Seedling Photomorphogenesis. The Plant Cell 25, 1243–1257 (2013).

56. Perron, N., Kirst, M. & Chen, S. Bringing CAM photosynthesis to the table: paving the way for resilient and productive agricultural systems in a changing climate. Plant Commun 100772 (2023) doi:10.1016/j.xplc.2023.100772.

57. Cabanettes, F. & Klopp, C. D-GENIES: dot plot large genomes in an interactive, efficient and simple way. PeerJ 6, e4958 (2018).

58. Liu, D. et al. Perspectives on the basic and applied aspects of crassulacean acid metabolism (CAM) research. Plant Science 274, 394–401 (2018).

59. Tao, S. et al. Single-Cell Transcriptome and Network Analyses Unveil Key Transcription Factors Regulating Mesophyll Cell Development in Maize. Genes (Basel) 13, 374 (2022).

60. Huang, S. et al. Overexpression of Ginkgo BBX25 enhances salt tolerance in Transgenic Populus. Plant Physiology and Biochemistry 167, 946–954 (2021).

61. Mariac, C. High molecular weight DNA extraction from plant nuclei isolation optimised for long-read sequencing. (2020).

62. Schalamun, M. & Schwessinger, B. High molecular weight gDNA extraction after Mayjonade, et al. optimised for eucalyptus for nanopore sequencing. (2017).

63. Fang, G., Hammar, S. & Grumet, R. A quick and inexpensive method for removing polysaccharides from plant genomic DNA. Biotechniques 13, 52–54, 56 (1992).

64. Kolmogorov, M. et al. metaFlye: scalable long-read metagenome assembly using repeat graphs. Nat Methods 17, 1103–1110 (2020).

65. Walker, B. J. et al. Pilon: An Integrated Tool for Comprehensive Microbial Variant Detection and Genome Assembly Improvement. PLOS ONE 9, e112963 (2014).

66. Simão, F. A., Waterhouse, R. M., Ioannidis, P., Kriventseva, E. V. & Zdobnov, E. M. BUSCO: assessing genome assembly and annotation completeness with single-copy orthologs. Bioinformatics 31, 3210–3212 (2015).

67. Holt, C. & Yandell, M. MAKER2: an annotation pipeline and genome-database management tool for second-generation genome projects. BMC Bioinformatics 12, 491 (2011).

68. Chang, S., Puryear, J. & Cairney, J. A simple and efficient method for isolating RNA from pine trees. Plant Mol Biol Rep 11, 113–116 (1993).

69. Li, H. Minimap2: pairwise alignment for nucleotide sequences. Bioinformatics 34, 3094– 3100 (2018).

70. Conde, D. et al. A robust method of nuclei isolation for single-cell RNA sequencing of solid tissues from the plant genus Populus. PLOS ONE 16, e0251149 (2021).

