## Supplementary Figures for "Mesophyll-Specific Circadian Dynamics of CAM Induction in the Ice Plant Unveiled by Single-Cell Transcriptomics"

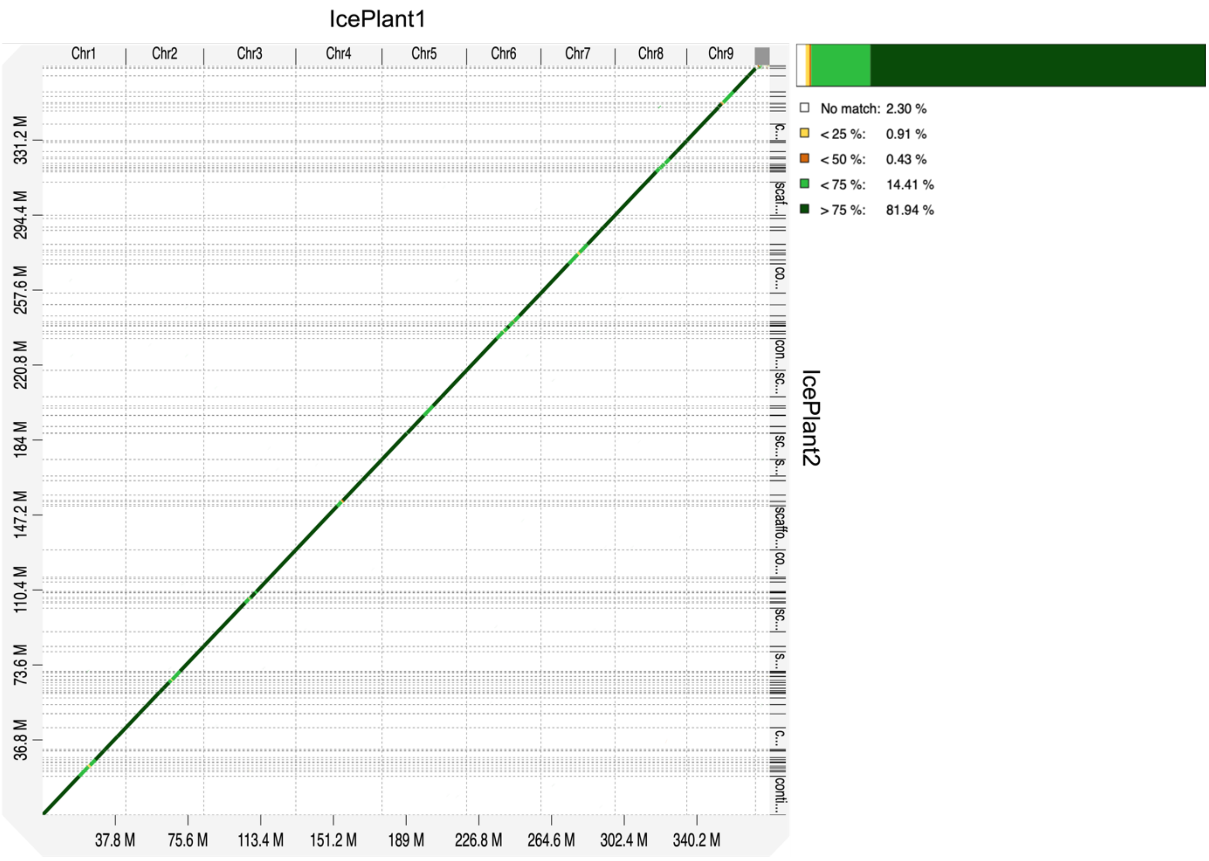


Supplementary Figure 1: Whole genome alignment of the IcePlant2 assembly from the present study (y-axis) against the previously released IcePlant1 assembly (x-axis) using D-Genies. The percentage values displayed in this alignment represent the similarity profile between the two genomes. This is calculated as the sum of match projections on the reference genome (here, IcePlant1) for each similarity category, divided by the total length of the reference genome.

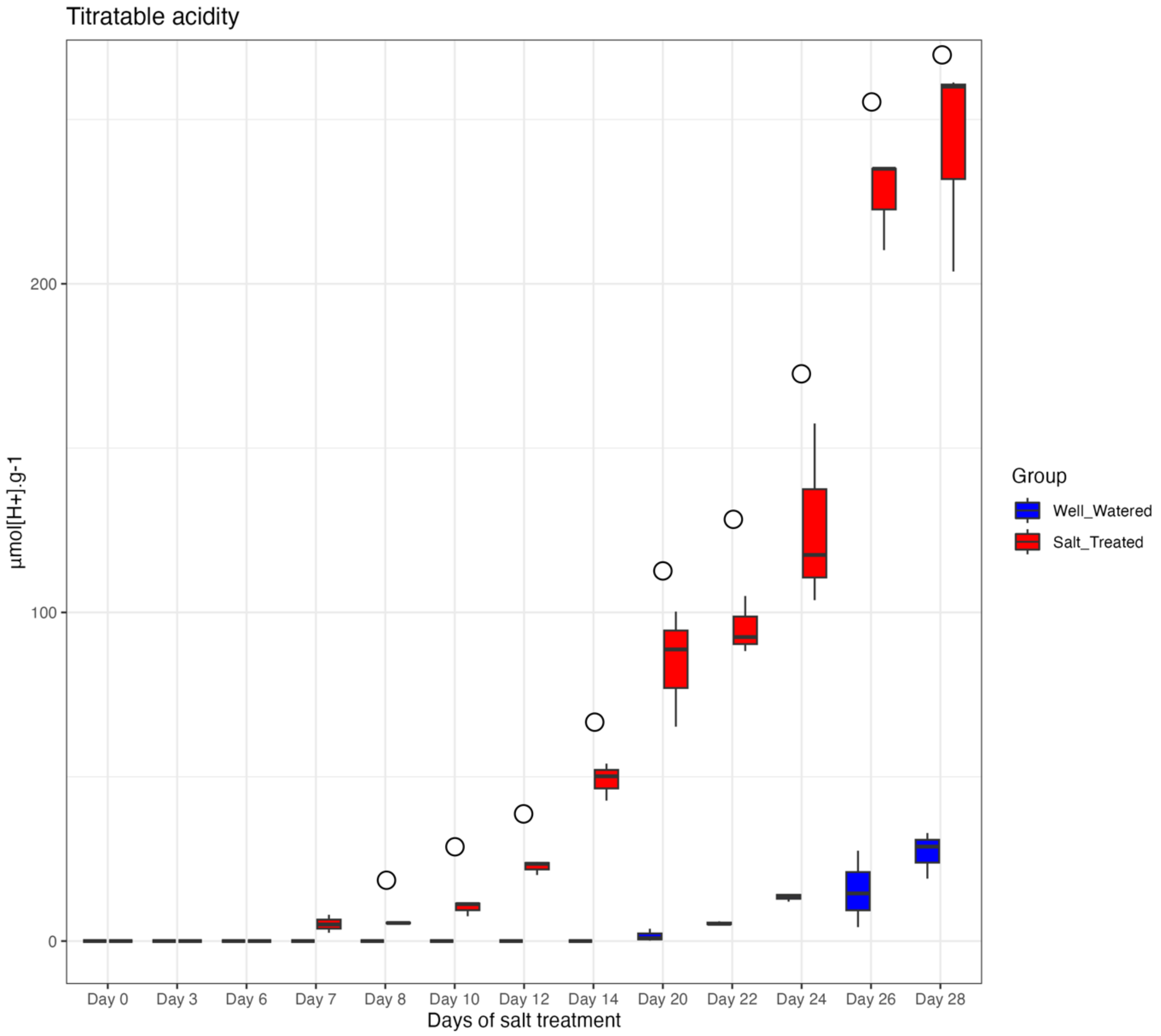


Supplementary Figure 2: Boxplot representation of the evolution of titratable acidity (measured in µmol[$H^+$].$g^-1$) in the leaves of well-watered and salt-treated ice plants. For each time-point, titration was performed on leaves from three biological replicates for each condition. Significant differences between the groups are marked by open circles, as determined by a paired t-test (p-value < 0.05). Each boxplot illustrates the median value (black line), the first and third quartiles (the lower and upper hinges), and the extremes of the data range (the top and bottom whiskers). No outliers are depicted in this graph.
